# Monocarboxylate transporters facilitate succinate uptake into brown adipocytes

**DOI:** 10.1101/2023.03.01.530625

**Authors:** Anita Reddy, Sally Winther, Nhien Tran, Haopeng Xiao, Josefine Jakob, Ryan Garrity, Arianne Smith, Evanna L. Mills, Edward T. Chouchani

**Affiliations:** Department of Cancer Biology, Dana-Farber Cancer Institute, Boston, MA, USA; Department of Cell Biology, Harvard Medical School, Boston, MA, USA; Novo Nordisk Foundation Center for Basic Metabolic Research, Faculty of Health and Medical Sciences University of Copenhagen, Copenhagen, Denmark; Department of Cancer Immunology and Virology, Dana-Farber Cancer Institute, Boston, MA, USA; Department of Immunology, Department of Immunology, Harvard Medical School, Boston, MA, USA

## Abstract

Uptake of circulating succinate by brown adipose tissue (BAT) and beige fat elevates whole body energy expenditure, counteracts obesity, and antagonizes systemic tissue inflammation in mice. The plasma membrane transporters that facilitate succinate uptake in these adipocytes remain undefined. Here we elucidate a mechanism underlying succinate import into BAT via monocarboxylate transporters (MCTs). We show that succinate transport is strongly dependent on the proportion of it present in the monocarboxylate form. MCTs facilitate monocarboxylate succinate uptake, which is promoted by alkalinization of the cytosol driven by adrenoreceptor stimulation. In brown adipocytes, we show that MCT1 primarily facilitates succinate import, however other members of the MCT family can partially compensate and fulfill this role in the absence of MCT1. In mice, we show that acute pharmacological inhibition of MCT1 and 2 decreases succinate uptake into BAT. Conversely, congenital genetic depletion of MCT1 alone has little effect on BAT succinate uptake, indicative of additional transport mechanisms with high capacity *in vivo*. In sum, we define a mechanism of succinate uptake in BAT that underlies its protective activity in mouse models of metabolic disease.

## Introduction

Beyond its role as a tricarboxylic acid (TCA) cycle intermediate, succinate is now known to regulate several physiological and pathophysiological processes (Mills & O’Neill, 2014; Murphy & Chouchani, 2022; Murphy & O’Neill, 2018). Succinate elicits biological adaptation via its accumulation and subsequent modulation of intracellular processes (Murphy & Chouchani, 2022). One consequence of accumulated succinate arises from its capacity to contribute reducing power to the mitochondrial respiratory chain, and in doing so to regulate the production of superoxide and hydrogen peroxide (Murphy, 2009). Accumulated succinate is also readily released from mitochondria and can regulate cellular adaptation through inhibition of alpha-ketoglutarate (αKG) dependent dioxygenases, a large family of enzymes with broad roles ranging from oxygen sensing, fatty acid catabolism, and regulation of the epigenome (Losman et al., 2020).

Additionally, accumulated succinate is now well appreciated to regulate physiology through its secretion and ligation of the G-protein coupled receptor, succinate receptor 1 (SUCNR1)(Aguiar et al., 2014; Gilissen et al., 2016; McCreath et al., 2015; Mills et al., 2021; Reddy et al., 2020; Sadagopan et al., 2007; van Diepen et al., 2017). Recent studies have shown that succinate is selectively released from cells via the plasma membrane monocarboxylate transporter (MCT) 1 (Andrienko et al., 2017; Bisbach et al., 2022; Prag et al., 2021; Reddy et al., 2020). This process is initiated by intracellular acidification, for example in response to acute metabolic perturbations such as muscle exercise or tissue hypoxia (Prag et al., 2021; Reddy et al., 2020). Acute acidification causes a portion of the intracellular succinate pool to become protonated, converting the metabolite from a dicarboxylate to a monocarboxylate, which renders it a transport substrate for MCT1. Following secretion, succinate signaling via SUCNR1 has been shown to regulate numerous physiological processes, including increased innervation following exercise training and cytokine production by macrophages (Aguiar et al., 2014; Littlewood-Evans et al., 2016; Mills et al., 2021; Reddy et al., 2020; Rubic et al., 2008; Sapieha et al., 2008; van Diepen et al., 2017; Vargas et al., 2009; Wu et al., 2020).

A major factor that antagonizes SUCNR1 signaling by succinate is the rate of succinate sequestration by tissues following its release. SUCNR1 activation occurs at a half-maximum effective concentration (EC50) between 28–56 μM, with ~99% responses achieved at 200 μM (He et al., 2004). Extracellular succinate readily achieves these concentrations following release (Correa et al., 2007; Hochachka & Dressendorfer, 1976; Mills et al., 2021; Reddy et al., 2020; Sadagopan et al., 2007). Also, depending on the physiological context, extracellular succinate levels can rapidly renormalize, or remain chronically elevated (Hochachka & Dressendorfer, 1976; Mills et al., 2021; Reddy et al., 2020; Sadagopan et al., 2007). The varied kinetics and fates of circulating succinate coincide with distinct physiological outcomes. For example, pulsatile elevation of extracellular succinate is associated with physiological remodeling of muscle tissue following exercise, while chronic elevation of extracellular succinate is linked to pro-fibrotic and pro-inflammatory signaling (Hochachka & Dressendorfer, 1976; Mills et al., 2021; Osuna-Prieto et al., 2021; Reddy et al., 2020; Sadagopan et al., 2007).

Brown adipose tissue (BAT) can sequester extracellular succinate (Mills et al., 2021; Mills et al., 2018). Uptake of succinate by BAT is linked to protection against various aspects of metabolic disease. Succinate uptake by BAT promotes uncoupling protein 1 (UCP1)-dependent thermogenesis, energy expenditure, and protection against obesity and glucose intolerance in mice (Mills et al., 2018). Additionally, sequestration of extracellular succinate by BAT antagonizes chronic SUCNR1 mediated liver inflammation and fibrosis in the context of non-alcoholic fatty liver disease (NAFLD) (Mills et al., 2021). While succinate uptake into BAT results in beneficial systemic effects, little is known about the mechanism of succinate transport in BAT or how it is regulated. Here we identify a key role for MCTs and pH-gradient facilitated succinate uptake.

## Materials and Methods

### Mouse Lines

Animal experiments were performed according to procedures approved by the Institutional Animal Care and Use Committee of the Beth Israel Deaconess Medical Center. Unless otherwise stated, mice used were male C57BL/6J (10 ± 2 weeks of age; Jackson Laboratories) and housed in a temperature-controlled (23°C) room on a 12-h light-dark cycle. Both male and female MCT1 KO mice were used. MCT1 flox mice were a gift from the Rothstein lab (Jha et al., 2020).

### Primary brown adipocyte isolation

Primary brown adipocytes were obtained and handled as previously described (Kir et al., 2014; Mills et al., 2018). Briefly, intrascapular brown adipose tissue was dissected from 2-to 10-day-old pups, washed in PBS and minced. The minced tissue was digested for 45 min at 37°C in PBS buffer containing 1.5 mg ml^-1^ collagenase B, 123 mM NaCl, 5 mM KCl, 1.3 mM CaCl_2_, 5 mM glucose, 100 mM HEPES, and 4% essentially fatty-acid-free BSA. The tissue suspension was filtered first through a 100 μm and then a 40 μm cell strainer and spun at 600 g for 5 min inbetween to pellet the stromal vascular fraction. The cell pellet was resuspended in adipocyte culture medium, plated, and maintained at 37°C in 10% CO_2_. Primary brown pre-adipocytes were counted and diluted to a concentration of 150,000 cell/ml and plated in the evening 12h before differentiation start. Plating was scaled according to surface area, for 24-well plates 0.5 ml (75,000 cells) were used/well. Brown preadipocytes were induced to differentiate with an adipogenic cocktail diluted in adipocyte culture medium (1 μM rosiglitazone, 0.5 mM IBMX, 5 μM dexamethasone, 0.114 μg ml^-1^ insulin, 1 nM T3, and 125 μM indomethacin). Two days after induction, and subsequently every 48hrs, cells were re-fed adipocyte culture medium containing 1 μM rosiglitazone, 1 nM T3, and 0.5 μg ml^-1^ insulin. Cells were fully differentiated 7 days after induction.

### ^14^C_4_-Succinate uptake assay

For radioactive uptake assays cells were grown in 24-well plates. On day of experiment tissue culture media was aspirated and the cells were washed once in pre-warmed (37°C) PBS. The cells were then preincubated for 30 min in assay buffer (DPBS with MgCl_2_ & CaCl_2_ supplemented with 2% BSA and 1 mM glucose; pH 7.5) at 37°C. For MCT inhibitor experiments the inhibitors were added at the specified concentrations in the preincubation buffer. After preincubation buffer was removed and replaced with 200 μl assay buffer containing [1,4-^14^C] succinic acid (American Radiolabeled chemicals) for 15 min (unless otherwise stated) at 37°C. pH of the assay buffer was adjusted with NaOH to pH 6.5, unless otherwise stated. The radioactive assay buffer contained 0.5 μCi/ml ^14^C-succinate and was supplemented with sodium succinate to a final concentration of 5 mM (unless otherwise stated). For metabolite competitor studies the indicated metabolite was added to the uptake buffer together with succinate. At the end of the 15 min cells were placed on ice and washed 3x with ice-cold PBS. Cells were lysed by pipetting in 200 μl 1% SDS, and 150 μl was transferred to a scintillation vial (Wheaton) containing 6 ml Ultima Gold scintillation cocktail (Perkin Elmer). Radiation in each sample was measured as counts/minute (CPM) using a TRI-CARB 2900TR scintillation counter. For each experiment at blank sample was run to determine background counts, and this value was subtracted from all experimental readings. For normalization total protein concentration was measured on an aliquot of the cell lysate with the Pierce BCA protein kit (Thermo Fisher) following the manufacturer’s instructions. Radioactive CPM was converted to succinate concentrations using the specific activity (55 mCi/ml) of ^14^C-succinic acid. The uptake was then normalized to total protein amount for each sample. (For experiments with preadipocytes and cell lines uptake data was normalized to cell counts instead of protein concentration).

### *In vitro* ^13^C_4_-Succinate uptake assay

For ^13^C_4_-succinate uptake assays, cells were grown in 24-well plates. On the day of the experiment, serum-free media (Gibco) containing ^13^C_4_-succinate (Cambridge Isotopes) was prepared and pH was adjusted to pH 7.4 using NaOH. For norepinephrine (NE) stimulation experiments, a specified amount of NE (Sigma Aldrich) was added to the ^13^C_4_-succinate containing media just prior to incubation. For MCT1 inhibitor experiments, cells were pre-incubated with AZD-3965 for 30 min. After pre-incubation, ^13^C_4_-succinate containing media was added and cells were incubated for specified amount of time. Media was aspirated and cells were washed twice on dry ice with icecold PBS. 100 μL of metabolite extraction buffer consisting of 80% HPLC MeOH containing D4-thymine, D4-glycocholate, and 15N4-inosine was added to each well. Cells were scraped and lysates were collected for further LC/MS analysis.

### Cytosolic pH measurements

Intracellular pH was measured using the ratiometric pH dye BCECF-AM (Invitrogen) (Khacho et al., 2014). Preadipocytes were plated in a 12-well plate and differentiated. On day 7, mature adipocytes were loaded with dye (1 mM) and incubated for 30 min at 37°C. Cells were imaged using a dual-excitation ratio of 480 nm and 440 nm and a fixed emission at 535 nm. A baseline image was obtained before norepinephrine (Sigma Aldrich) was administered. A calibration curve was created using nigericin and high K^+^.

### siRNA mediated knockdown in brown adipocytes

Forward and reverse siRNA transfections were performed as previously described (Isidor et al., 2016). Transfections were carried out on day 4 of differentiation and experiments were conducted on day 7. For forward transfections, pre-adipocytes were plated in a 24-well plate at a concentration of 50,000 cells/well. For reverse transfections, day 4 differentiated brown adipocytes were plated at a concentration of 475,000 cells/well in a 24-well plate. Final concentrations of siRNA and lipofectamine RNAiMAX was 90 nM and 9 μl/ml, respectively. siRNAs (Sigma-Aldrich) used were: Mct1 (SASI_Mm01_00112354), Mct2 (SASI_Mm01_00096521), Mct4 (SASI_Mm01_00119743), Slc13a3 (SASI_Mm01_00057701). MISSION^®^ siRNA Universal Negative Control #1 (Sigma-Aldrich, SIC001) was used as control siRNA.

### Sample preparation for metabolite analysis

Following intervention, 100 uL of ice-cold extraction buffer consisting of 80% methanol containing D4-thymine, D4-glycocholate, and 15N4-inosine was added to each well of a 24-well plate. Cells were immediately harvested by scraping on dry ice and transferred to a 1.5 mL microcentrifuge tube. Lysates were centrifuged twice at max speed (10 min, 21,100 g, 4°C) and supernatant was collected. For *in vivo* experiments, mice were sacrificed by cervical dislocation following intervention and tissues were quickly harvested and immediately snap-frozen in liquid nitrogen. Samples were then weighed and homogenized with extraction buffer in a 4:1 (μL extraction buffer: mg of tissue) ratio. Tissues were centrifuged twice (10 min, 21,100 g, 4°C) and supernatant was collected. Supernatant was further diluted 1:10 (sample:80% methanol) with 80% HPLC grade methanol. For plasma collection, blood was collected via cheek bleed into a heparin column (Becton Dickson), centrifuged (10 min, 1,000 g, 4°C), and immediately snap-frozen. Metabolites were then extracted by adding extraction buffer in a 4:1 (μL of extraction buffer: μL of plasma) ratio. Extracted metabolites were further diluted 1:10 (sample:80% methanol) with 80% HPLC grade methanol.

### LC/MS metabolite analysis

Metabolite separation was performed using a Luna-HILIC column (Phenomenex) on an UltiMate-3000 TPLRS LC. A linear 10-min gradient of 10% mobile phase A (20 mM ammonium acetate and 20 mM ammonium hydroxide in 95% water and 5% ACN) and 90% mobile phase B (10 mM ammonium hydroxide in 75:25 v/v ACN/MeOH) to 99% mobile phase A was used, and metabolites were analyzed by a Q-Exactive^™^ HF-X mass spectrometer (ThermoFisher). Negative ion mode was used with full scan analysis over m/z 70-750 m/z at 60,000 resolution, 1e6 AGC, and 100 ms maximum ion accumulation time. Ion spray voltage was set at 3.8 kV, capillary temperature was at 350°C, probe heater temperature was at 320 °C, sheath gas flow was set at 50, auxiliary gas was set at 15, and S-lens RF level was set at 40. TraceFinder (ThermoFisher) was used to annotate and quantify the metabolite peaks.

### *In vivo* ^13^C_4_-Succinate tracing

Mice were housed at thermoneutrality (29°C) for one week prior to experiment. Mice were then injected with [U^13^C] succinate (10 mg/kg, 2 min) (Cambridge Isotope Laboratories) via intravenous tail vein. Intravenous injections were performed as a bolus over 20 s. Mice were sacrificed by cervical dislocation. Tissues and plasma were harvested and immediately snap frozen in liquid nitrogen and stored at −80°C for further analysis.

### *In vivo* chemical inhibition of MCTs

Mice were injected intraperitoneally with AZD 5 mg/kg, 2 h prior to intravenous tail vein injection with [U^13^C] succinate (10 mg/kg, 2 min) (Cambridge Isotope Laboratories). Intravenous injections were performed as a bolus over 20 s. Tissues were extracted and snap frozen in liquid nitrogen and stored at −80°C until MS analysis was performed.

### Gene expression analysis (qPCR)

Total RNA from cellular siRNA experiments were extracted using TRIzol (Invitrogen) and purified using phenol:chloroform extraction with isopropanol precipitation. Purified RNA was quantified using a Nanodrop 2000 UV-visible spectrophotometer. cDNA was prepared according to manufacturer’s instructions from 2 ug total RNA using a high-capacity cDNA reverse transcription kit (Applied Biosystems). qPCR reactions were performed in a 384-well format using GoTaq qPCR Master Mix (Promega) and run on an ABI PRISM 7900HT real time PCR system (Applied Biosystems). Relative gene expression was calculated by the delta delta Ct method using mouse cyclophilin A expression as endogenous control. All fold changes are expressed normalized to the universal negative siRNA control. Primer pairs are as follows: *Cyclophilin A*, FW 5’-GGAGATGGCACAGGAGGAA-3’, RV 5’-GCCCGTAGTGCTTCAGCTT-3’; *Mct1*, FW 5’-GTGCAACGACCAGTGAAGTA-3’, RV 5’-ACAACCACCAGCGATCATT-3’; *Mct2*, FW 5’-GTTGGCTTCAATGGAGGTTT-3’, RV 5’-AAAACAGGACTTCCAGCCAT-3’; *Mct4*, FW 5’-TCCTGCTGGCTATGCTCTA-3’, RV 5’-CAAAGAGGCCACCCACAA-3’; *Slc13a3*, FW 5’-AGCTCAAGAGTTTCTTCCCA-3’, RV 5’-CGTCAGCTCGTATCTTGGAT-3’.

### Brown adipose tissue proteomics

BAT samples were lysed in the lysis buffer (100 mM 4-(2-hydroxyethyl)-1-piperazineethanesulfonic acid (HEPES) pH 8.5, 8 M urea, 2% SDS, 1p/15 mL Roche cOmplete^™^ protease inhibitors) to 1 mg/mL protein based on BCA assay results. Disulfides were reduction with 5 mM tris(2-carboxyethyl)phosphine (TCEP) at 37 °C for 1 h, followed by alkylation with 25 mM iodoacetamide for 25 min at room temperature in the dark. Protein precipitation was performed by the methanol-chloroform method (Wessel & Flugge, 1984), and proteins were digested resuspended in 200 mM N-(2-Hydroxyethyl)piperazine-N’-(3-propanesulfonic acid) (EPPS) buffer pH=8, using a combination of Lys-C and trypsin at an enzyme-to-protein ratio of 1:100 overnight at 37 °C, followed by an additional 4 h digestion with trypsin 1:100. Samples were then subjected to a microBCA measurement for peptide quantification, and 25 μg peptides from each sample were labeled by TMTpro-16 reagents (Li et al., 2020) for 1 h at room temperature following the streamlined-TMT protocol (Navarrete-Perea et al., 2018). The reaction was quenched using 2 μl of 5% hydroxylamine for 15 min. A ratio-check was performed by mixing 2 μL of peptides from each channel, desalted via StageTip, and analyzed by LC-MS. Samples were combined according to the ratio check to ensure equal loading of proteins from each sample, then desalted with Waters SepPak cartridges and dried. Peptide were fractionated using high-pH HPLC. Each fraction was then desalted via StageTip, dried in a speedvac, and reconstituted in a solution containing 5% ACN and 5% FA for liquid chromatography tandem mass spectrometry (LC-MS/MS).

### LC-MS/MS

2 μg of peptides were loaded onto an in-house 100-μm capillary column packed with 35 cm of Accucore 150 resin (2.6 μm,150 Å). An Orbitrap Eclipse Tribrid Mass Spectrometer (Thermo) coupled with an Easy-nLC 1200 (Thermo) and FAIMSPro (Thermo) were used for protein measurements. Peptides were separated using a 180-min gradient consisting of 2% - 23% ACN, 0.125% FA at 500 nl/min flow rate. Field asymmetric waveform ion mobility spectrometry (FAIMS) separation of precursors (Schweppe et al., 2019) were carried out with default settings and multiple compensation voltages (−40V/-60V/-80V). Peptide ions were collected in data-dependent mode using a mass range of m/z 400-1600 using 2 s cycles and 120,000 resolution. Singly-charged ions were discarded, and multiply-charged ions were selected and subjected to fragmentation with standard automatic gain control (AGC) and 35% normalized collisional energy (NCE) for MS2, with a dynamic exclusion window of 120 s and maximum ion injection time of 50 ms. Quantification of TMT reporter ion were performed using the multinotch SPS-MS3 method (McAlister et al., 2014) with 45% NCE for MS3.

### Database searching and protein quantification

Raw files were searched using the Comet algorithm (Eng et al., 2013) on Masspike reported previously (Huttlin et al., 2010). Database searching included all mouse (Mus musculus) entries from UniProt (http://www.uniprot.org, downloaded July 29^th^, 2020) and the reversed sequences as well as common contaminants (keratins, trypsin, etc). Peptides were searched using the following parameters: 25 ppm precursor mass tolerance; 1.0 Da product ion mass tolerance; fully tryptic digestion; up to three missed cleavages; variable modification: oxidation of methionine (+15.9949); static modifications: TMTpro (+304.2071) on lysine and peptide N terminus, carboxyamidomethylation (+57.0214637236) on cysteines. The target-decoy method was used for false discovery rate (FDR) control (Elias & Gygi, 2007; Huttlin et al., 2010; Peng et al., 2003). FDR was < 1% on peptide level for each MS run and peptides that are shorter than seven amino acids were not used. Proteins were assembled to <1% protein-level FDR. TMT reporter ions were used for quantification of peptide abundance. Peptides with summed signal-to-noise (S/N) lower than 160 across 16 channels or isolation specificity lower than 70% were discarded. Normalization was performed to ensure equal protein loadings in every sample. Proteins were quantified by summing peptide TMT S/N.

## Results

### Brown adipocytes import extracellular succinate in a pH dependent manner

We first used ^14^C_4_-succinate to characterize properties of the succinate transport system in differentiated mouse primary brown adipocytes. As expected, externally applied succinate resulted in concentration-dependent transport across the range of 0.01–20 mM extracellular succinate (**Figure 1a**). Succinate transport occurred at a maximal rate of 800 pmol/mg protein/min (**Figure 1a**). We next monitored the fate of internalized succinate by tracking uptake and metabolism of ^13^C_4_-succinate using mass spectrometry (**Figure 1b**). At external concentrations as low as 10 μM, uptake and mitochondrial metabolism of extracellularly applied succinate occurred rapidly, with ^13^C_4_-fumarate and ^13^C_4_-malate accumulating within cells as early as two minutes following application of ^13^C_4_-succinate (**Figure 1c**).

**1.**
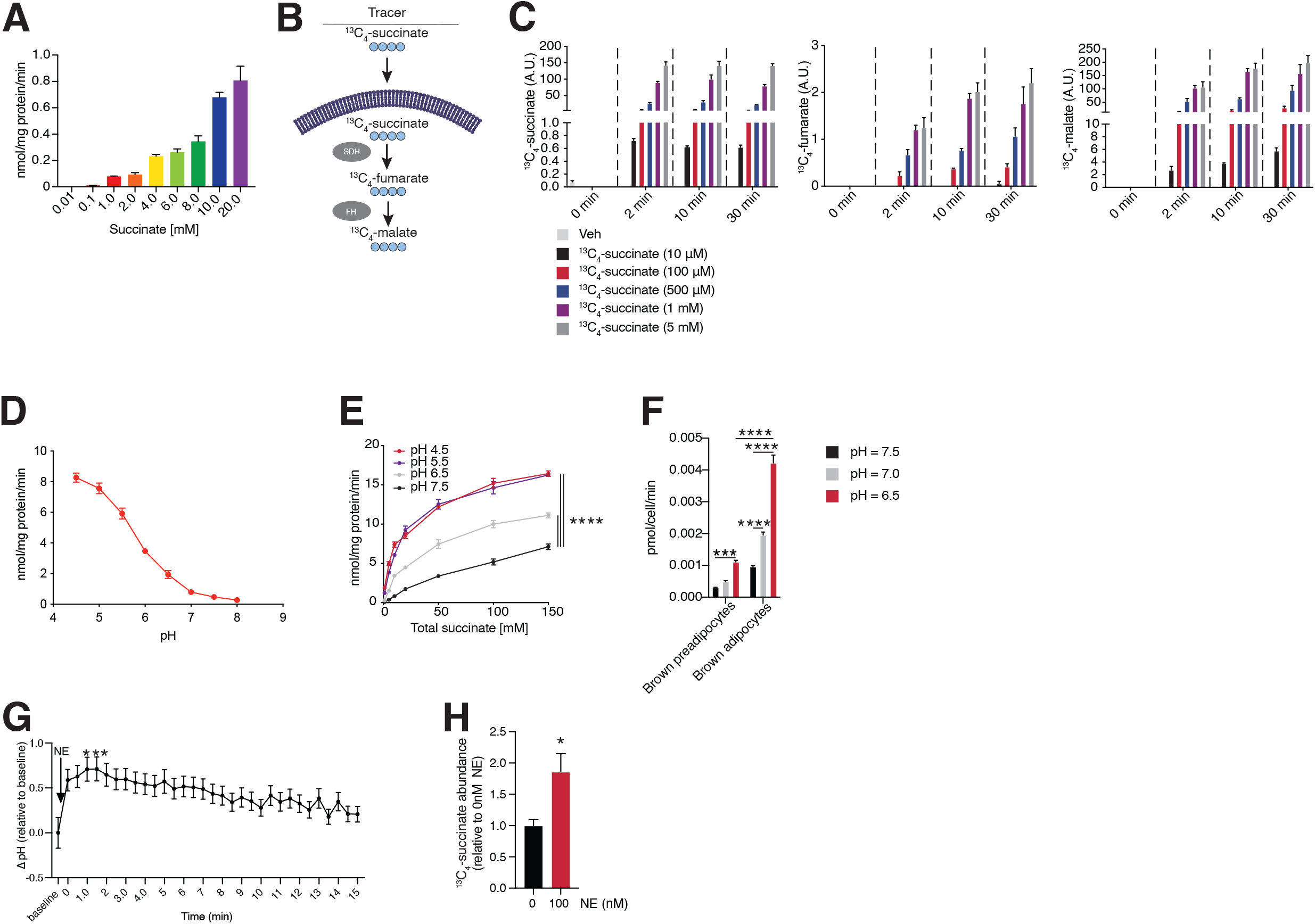
Primary brown adipocytes transport extracellular succinate in a pH dependent manner. a. Brown adipocytes were treated with ^14^C_4_-succinate at the indicated concentrations (pH = 7.45) for 15 min and ^14^C_4_-succinate uptake was monitored (n = 3). b. Schematic depicting transport and metabolism of ^13^C_4_-succinate tracer. c. Brown adipocytes were treated with ^13^C_4_-succinate at indicated concentrations and times (pH 7.45). Intracellular, relative abundance ^13^C_4_-succinate and its downstream metabolites, ^13^C_4_-fumarate and ^13^C_4_-malate, was assessed by LC/MS analysis (n = 4). d. Media pH was adjusted to indicated values and brown adipocytes were treated with ^14^C_4_-succinate (5 mM) for 15 min. ^14^C_4_-succinate uptake was monitored (n = 3). Brown adipocyte protein concentration was determined by bicinchoninic acid (BCA) assay. e. Brown adipocytes were treated with ^14^C_4_-succinate at indicated concentrations for 15 min under different media pH conditions, as shown. ^14^C_4_-succinate uptake was monitored (n = 3). Brown adipocyte protein concentration was determined by BCA assay. f. Primary brown adipocytes and preadipocytes were treated with ^14^C_4_-succinate (5mM) for 15 min under various media pH conditions, as indicated. ^14^C_4_-succinate uptake was monitored (n = 3). g. Brown adipocytes were loaded with the ratiometric, pH-sensitive dye, BCECF-AM (1 mM) for 30 min. Cells were then treated with 1 μM norepinephrine (NE). Images were collected every 30 s for the duration of 15 min using a dual-excitation ratio of 480 nm and 440 nm and a fixed emission at 535 nm. A baseline image, before NE administration, was also collected. Intracellular ΔpH of brown adipocytes was determined by using a calibration curve. h. Brown adipocytes were treated with 100 μM ^13^C_4_-succinate in the presence or absence of 100 nM NE for 2 min. Relative abundance of intracellular ^13^C_4_-succinate was determined by LC/MS analysis (n = 6). Data presented as mean ± SEM. One-way ANOVA test was performed for multiple comparisons involving one independent variable (1g), two-way ANOVA test was performed for comparisons involving two independent variables (1e, 1f), and two-tailed Student’s t test for pairwise comparisons (1h). *p ≤ 0.05, **p ≤ 0.01, ***p ≤ 0.001, ****p ≤ 0.0001.

Previous work performed in muscle, myotubes, and the ischemic heart showed that decreasing intracellular pH promotes succinate export (Prag et al., 2021; Reddy et al., 2020). On this basis, we examined whether succinate uptake into brown adipocytes could be stimulated by decreasing extracellular pH **(Figure 1d)**. We observed an inverse relationship between extracellular pH and the amount of succinate transported: as extracellular pH lowered, brown adipocytes imported more succinate (**Figure 1d**). Increased uptake was attributable to elevation in initial linear rate of transport of succinate as extracellular pH lowered (**Figure 1e**). Next, we monitored whether capacity for brown adipocyte succinate uptake depended on differentiation state. Using primary brown adipocytes, we assessed the ability of pre-adipocytes to transport succinate. Pre-adipocytes exhibited significantly reduced capacity for succinate uptake compared to differentiated brown adipocytes (**Figure 1f**). Previous work showed BAT preferentially accumulates succinate upon cold exposure (Mills et al., 2018). Adrenergic signaling stimulated by norepinephrine (NE) release from sympathetic nerves in BAT is a key signaling event that occurs in response to cold (Kawate et al., 1994; Wang et al., 2011; Young et al., 1982). Moreover, previous work has demonstrated cytosolic alkalinization following activation of β-adrenergic signaling (Bast-Habersbrunner & Fromme, 2020; Chinet et al., 1978; Lee et al., 1994). In line with this we observed that stimulation of brown adipocytes with NE resulted in a rapid alkalinization of cytosolic pH by ~0.6 pH units. (**Figure 1g**). Based on the ΔpH dependence of succinate export (Prag et al., 2021; Reddy et al., 2020), we hypothesized that a ΔpH imposed by cytoplasmic alkalinization induced by adrenergic signaling could promote succinate uptake by brown adipocytes. In support of this hypothesis, we observed that upon NE treatment uptake of succinate into brown adipocytes was significantly potentiated (**Figure 1h**). Taken together these findings indicate that succinate transport occurs at physiological pH but can be enhanced by a ΔpH gradient across the plasma membrane.

### Monocarboxylate transporters (MCTs) facilitate succinate uptake in brown adipocytes

We next sought to determine the protein(s) responsible for mediating succinate uptake in brown adipocytes. We first assessed a role for the canonical transporter of succinate, solute carrier family 13 member 3 (SLC13A3). Selective depletion of SLC13A3 in brown adipocytes had no effect on succinate uptake (**Figure 2a**). We next considered the fact that succinate uptake in brown adipocytes is pH dependent. Succinate is a dicarboxylate with a monocarboxylic pKa of 5.69 and a fully protonated pKa of 4.21 (Gokel & Dean, 2004). The monocarboxylic pKa of succinate is much higher than that of other TCA cycle intermediates rendering this metabolite amenable to protonation at physiological pH. As such, across the pH range 4.5 - 7.5 the proportion of succinate existing in the monocarboxylate form will vary substantially. We examined the relationship between brown adipocyte succinate uptake rate, extracellular pH, and the proportion of succinate existing as a monocarboxylate. This analysis demonstrated a strong relationship between the amount of succinate present in the monocarboxylate form and the rate of succinate sequestration by brown adipocytes (**Figure 2b,c**). Taken together, this data strongly suggests that the monocarboxylate form of succinate is the major transported form.

**2.**
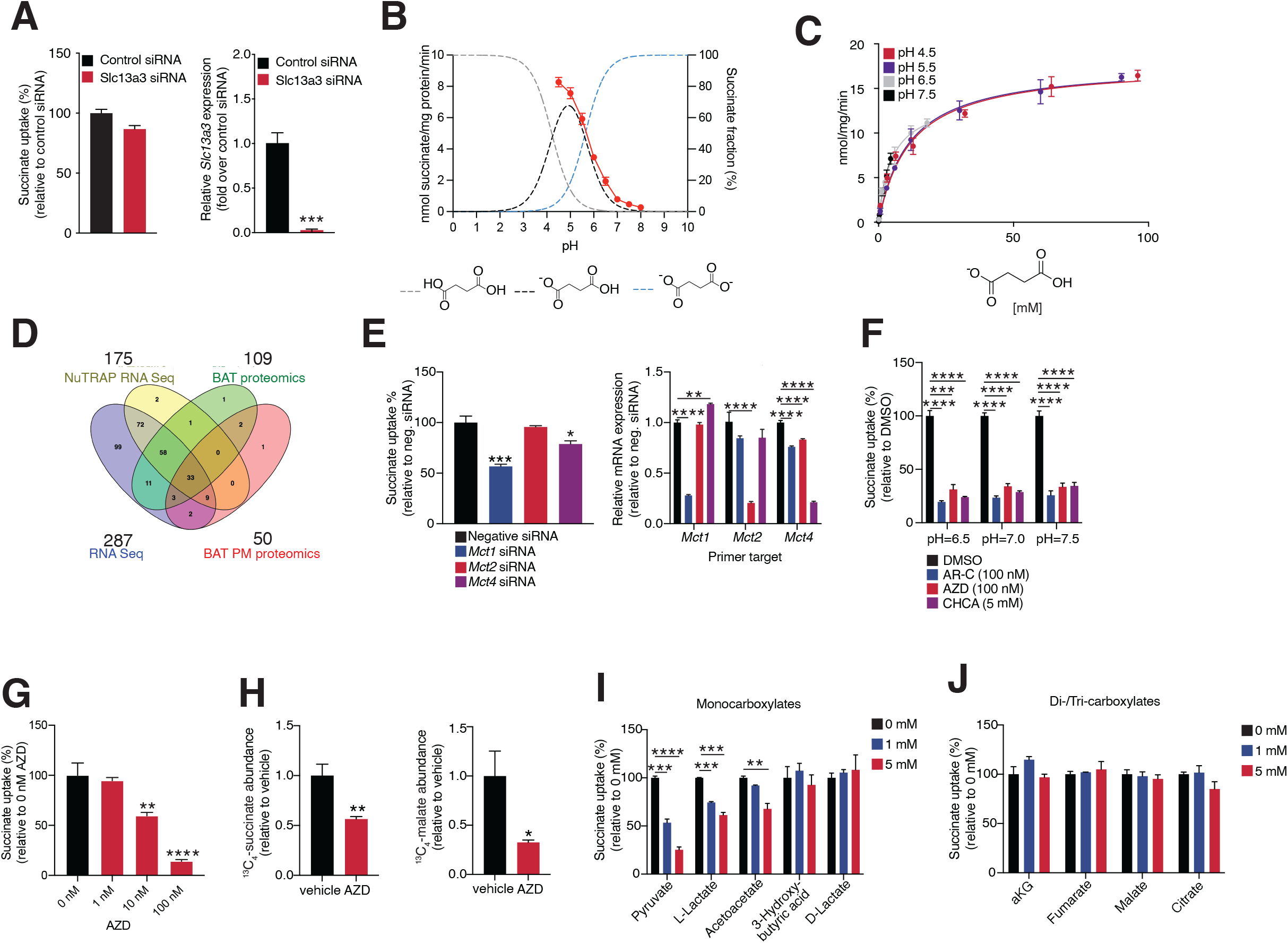
The MCT family of proteins are capable of transporting succinate *in vitro*. a. The plasma membrane, dicarboxylate transporter, *Slc13a3*, was genetically depleted in brown adipocytes. Knockdown was confirmed by qPCR. Cells were then treated with 5 mM ^14^C_4_-succinate (pH = 6.5) for 15 min and transport was monitored (n = 3). b. Schematic of figure 1D and succinate pKa curve. c. Rate of ^14^C_4_-succinate transport determined in figure 1E was recalculated to reflect the concentration of protonated ^14^C_4_-succinate present. d. Comparative screen to identify potential plasma membrane (PM) transporters of succinate present in brown adipose tissue (BAT). e. MCT proteins were genetically depleted from brown adipocytes. Knockdown was confirmed by qPCR. Cells were then treated with 5 mM ^14^C_4_-succinate (pH = 7.0) for 15 min and uptake was monitored (n = 3). f. Brown adipocytes were treated with MCT inhibitors AR-C155858, AZD3965, and CHCA at indicated concentrations. 5 mM of ^14^C_4_-succinate was administered to cells for 15 min under various pH conditions and uptake was monitored (n = 3). g. Brown adipocytes were treated with AZD3965 at concentrations indicated and 5 mM of ^14^C_4_-succinate for 15 min (pH = 6.5). ^14^C_4_-succinate transport was monitored (n = 3). h. Brown adipocytes were pre-treated with 100 nM of AZD3965 for 30 min and then were administered 10 μM ^13^C_4_-succinate for 2 min (pH = 7.45). Relative intracellular abundance of ^13^C_4_-succinate and downstream metabolite ^13^C_4_-malate was assessed by LC/MS analysis (n = 6). i-j. Competition assays were performed by treating brown adipocytes with 5 mM ^14^C_4_-succinate for 5 min in the presence of structurally similar metabolites, monocarboxylates (i) and di/tricarboxylates (j) at concentrations indicated (pH = 6.5). ^14^C_4_-succinate uptake was assessed (n = 3). All data presented as mean ± SEM. Two-tailed Student’s t-test was performed for pairwise comparisons (2a, 2h), and one-way ANOVA was performed for multiple comparisons involving one independent variable (2e, 2f, 2g, 2i, 2j) *p ≤ 0.05, **p ≤ 0.01, ***p ≤ 0.001, ****p ≤ 0.0001.

We next examined which small molecule carboxylate transporters are present in brown and beige adipocytes that could explain the basis for transport of monocarboxylic succinate across the plasma membrane. We compared four RNA and proteomic data sets comprising mouse brown/beige adipocytes to identify potential transporters of succinate (Roh et al., 2017). From this comparative analysis, MCT1 emerged among the common small molecule carboxylate transporters (**Figure 2d**). MCT1 belongs to the MCT family of proteins that include MCT1, 2, and 4. These transporters facilitate the diffusion of small monocarboxylates such as L-lactate and pyruvate across the plasma membrane. This transport is proton linked and is driven by changes in proton concentration (Halestrap & Price, 1999). Moreover, previous studies of muscle, heart, and recombinant protein have shown that MCT1 can transport succinate in its monocarboxylate form (Andrienko et al., 2017; Bisbach et al., 2022; Prag et al., 2021; Reddy et al., 2020).

Genetic depletion of MCT1 in brown adipocytes resulted in an attenuation in succinate uptake, whereas depletion of MCT4 had minimal effect and MCT2 no effect on uptake (**Figure 2e**). Chemical inhibition of the MCTs resulted in a similar attenuation of succinate transport (**Figure 2f**). AZD-3965, a potent inhibitor of MCT1 (IC_50_ = 1.6 nM) (Bola et al., 2014; Curtis et al., 2017) and at higher concentrations an inhibitor of MCT2 (IC_50_ = 20 nM) (Curtis et al., 2017), induced a concentration-dependent decrease in ^14^C_4_-succinate uptake (**Figure 2g**). Administering AZD-3965 at concentrations that inhibit MCT1 and MCT2 similarly resulted in a decrease in ^13^C_4_-succinate uptake and its downstream product of mitochondrial catabolism, ^13^C_4_-malate (**Figure 2h**). Extracellular co-application of known transport substrates of the MCT family of proteins also decreased succinate uptake by brown adipocytes, suggesting that these metabolites are competing for the same transporter as succinate (**Figure 2i**). Conversely, there was no effect on succinate uptake in the presence of excess di- or tricarboxylates (**Figure 2j**). Taken together, these data suggest MCT1, and to a lesser extent MCT2, can facilitate succinate uptake in brown adipocytes *in vitro*.

### Acute inhibition of MCT1 and MCT2 attenuates succinate uptake into mouse BAT *in vivo*

We next examined whether MCT1 regulates succinate uptake into mouse BAT *in vivo*. We generated an adipocyte-specific MCT1 knockout (KO) mouse by crossing a MCT1 flox mouse with a mouse expressing cre under the adiponectin promoter (**Figure 3a**). We confirmed ablation of MCT1 by performing quantitative proteomics on interscapular BAT from MCT1 KO and wildtype (WT) mice. We observed a significant decrease in MCT1 protein abundance along with its accessory protein basigin (BSG) (**Figure 3b**). This analysis provided no evidence for compensatory upregulation of other transporters in the fat-specific KO animals (**Figure 3c**). We next injected WT and MCT1 KO animals with ^13^C_4_-succinate and harvested interscapular BAT and subcutaneous adipose tissue (SAT) which contains both white and beige adipocytes (Garcia et al., 2016; Wu et al., 2012), and performed liquid chromatography/mass spectrometry (LC/MS) analysis to assess succinate uptake and metabolism in these tissues (**Figure 3d**). There was no significant difference in succinate uptake or metabolism in BAT (**Figure 3e**) or SAT (**Figure 3f**) from WT and adipocyte MCT1 KO.

**3.**
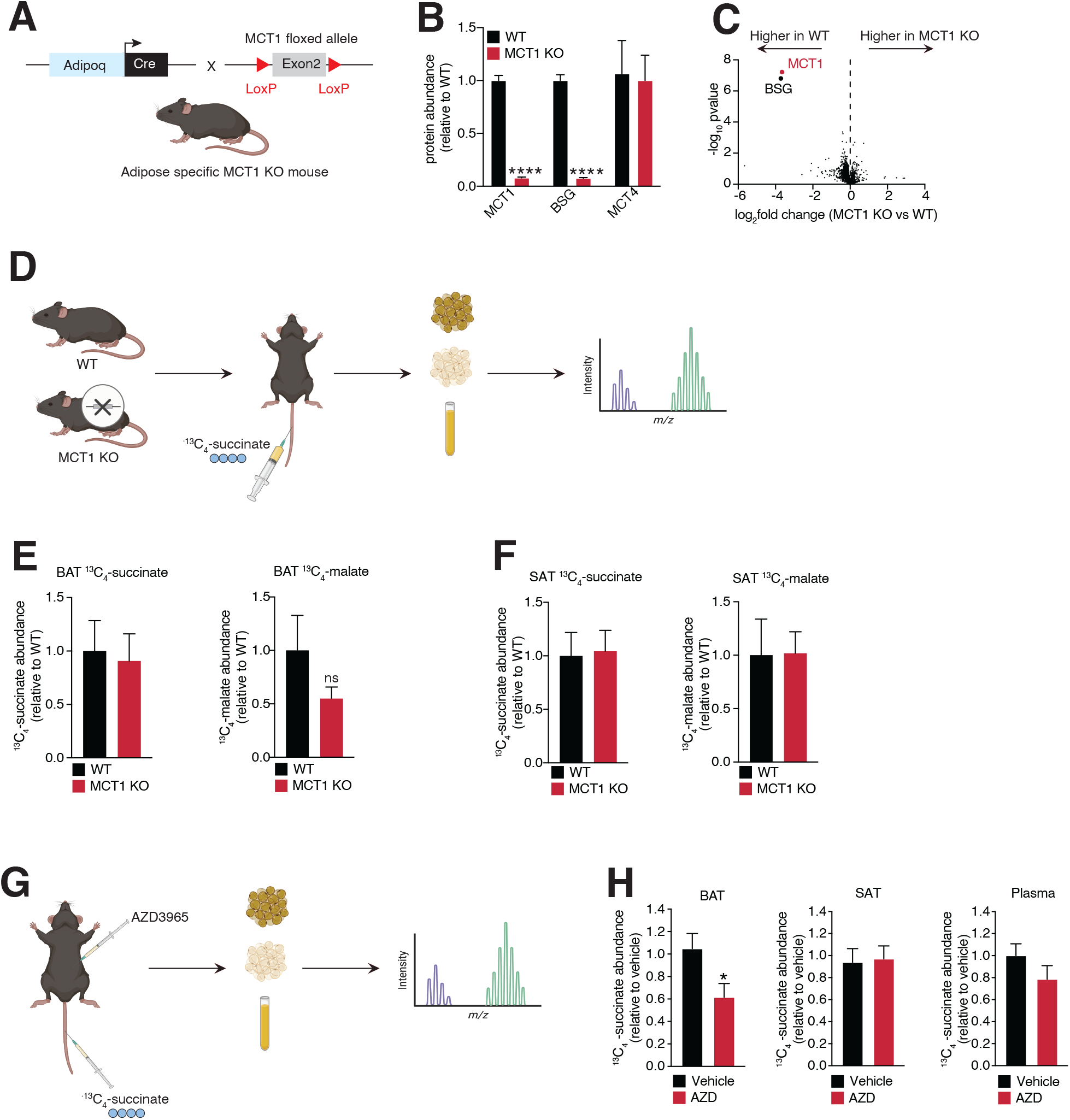
AZD3965 decreases succinate transport into BAT, *in vivo*. a. Schematic detailing the generation of the fat-specific MCT1 KO mouse. b. Proteomic analysis was performed on BAT harvested from wild-type (WT) and MCT1 KO mice to assess the relative abundance of MCT1, MCT4, and accessory protein basigin (BSG) (n = 4). c. Global proteomic analysis was performed on BAT harvested from WT and MCT1 KO mice to determine possible off-target changes in protein abundance in the absence of MCT1 (n = 4). d. Schematic outlining experimental procedure of *in vivo* ^13^C_4_-succinate tracing in WT and MCT1 KO mice. e-f. WT (n = 11) and MCT1 KO (n = 8) mice were housed at thermoneutrality (TN) for one week and were then administered ^13^C_4_-succinate (10 mg/kg) via tail-vein injection. 2 min following injection mice were sacrificed and BAT (e) and subcutaneous adipose tissue (SAT) (f) were harvested. Metabolites were extracted from these tissues and relative abundance of ^13^C_4_-succinate and ^13^C_4_-malate was determined. g. Schematic detailing experimental procedure of *in vivo* ^13^C_4_-succinate tracing following AZD3965 administration. h. WT mice were administered either 5 mg/kg AZD3965 (n = 10) or saline (n =16) via i.p. injection 2h prior to tail-vein injection of ^13^C_4_-succinate (10 mg/kg). 2 min following ^13^C_4_-succinate injection, BAT, SAT, and plasma were harvested. Metabolite extraction was performed on tissues and relative ^13^C_4_-succinate abundance was assessed via LC/MS analysis. All data presented as mean ± SEM. Two-tailed Student’s t-test was performed on pairwise comparisons (3b, 3e, 3f, 3h). *p ≤ 0.05, ****p ≤ 0.0001.

The discordance of our *in vivo* observations with those found in isolated brown adipocytes led us to consider possible explanations for these differences. One potential difference arose in the nature of acute versus chronic MCT depletion or inhibition. Our *in vitro* studies employed siRNA mediated knockdown (several days), or pharmacological inhibition (minutes). Conversely, our *in vivo* analysis was performed in mice with a congenital lack of MCT1, which could lead to altered development of thermogenic adipocytes and reliance on alternative modes of monocarboxylate transport. Moreover, the possibility of redundance in succinate uptake in BAT mediated by MCTs was supported by our *in vitro* analyses. Specifically, data from acute *in vitro* application of AZD-3965 suggests that both MCT1 and 2 are responsible for succinate transport since only higher concentrations (100 nM) led to maximal inhibition of succinate uptake (**Figure 2g**). This concentration would inhibit both MCT1 and MCT2, suggesting that MCT2 could facilitate succinate transport *in vivo*, in the absence of MCT1. To examine this possibility, we acutely inhibited MCT1 and 2 in WT mice using AZD-3965 and injected ^13^C_4_-succinate (**Figure 3g**). AZD-3965 treatment significantly decreased ^13^C_4_-succinate uptake by BAT but not SAT (**Figure 3h**). Taken together, these data indicate that MCTs facilitate succinate uptake into BAT *in vivo* and suggests a role for other MCTs in addition to MCT1.

## Discussion

Regulation of SUCNR1 by circulating succinate has emerged as a key node of metabolite control over tissue remodeling. The dynamics of succinate-SUCNR1 signaling appears to be a major component dictating the nature of the biological consequence of succinate action. Acute pulsatile elevation in extracellular succinate elicits distinct remodeling processes, when compared to chronic elevation. Additionally, SUCNR1 agonism appears to promote distinct adaptive responses depending on the SUCNR1-expressing cell type sensing extracellular succinate.

The dynamic nature of SUCNR1 agonism by succinate requires a better understanding of the mechanisms that control succinate secretion into, and sequestration from, extracellular fluids. Here we focused our attention on brown adipocytes, a cell type with a particular avidity for uptake of extracellular succinate. Brown adipocyte succinate uptake regulates thermogenic respiration in BAT and beige fat and whole-body energy expenditure. Moreover, succinate uptake by these cells has been shown to antagonize pathogenic SUCNR1 activation cause by chronically elevated circulating succinate. Here we show that brown adipocyte uptake of succinate depends on the proportion of it in the monocarboxylic form, and relies substantially on MCT family proteins, in particular MCT1. It is noteworthy that a mechanism through which succinate is acquired by brown adipocytes appears to be an inversion of the succinate export mechanism recently discovered in a variety of cell types (Bisbach et al., 2022; Prag et al., 2021; Reddy et al., 2020).

Studies in exercising muscle, ischemic heart, and mouse retinas have shown succinate export via MCT1 to be dependent on intracellular acidification occurring in response to changes in cellular metabolism. For example, during exercise, the combination of decreased oxygen tension and increased glycolytic flux results in acute intracellular acidification, with cytosolic pH falling as low as 6.4 (Robergs et al., 2004). Other physiological conditions, such as ischemia, can also lead to acute acidification of the cytosol (Prag et al., 2021). This pH regulation of succinate export positions succinate as an ideal signaling molecule since it can relay changes in bioenergetic status to the surrounding environment. Because the pKa of succinate is high compared to other mitochondrial dicarboxylates, a significant portion of the intracellular pool is readily protonated at physiological pH, converting the metabolite from a dicarboxylate to a monocarboxylate. Once protonated, succinate is then secreted from the cell via MCT1. The decrease in pH serves a dual purpose, not only does it increase the pool of monocarboxylate succinate, but it also supplies protons required for MCT-facilitated transport.

MCTs facilitate diffusion of carboxylates across the plasma membrane, and the rate and directionality are dictated by the concentration gradient of the carboxylate and protons. Notably, transport of the monocarboxylate form of succinate by MCTs would carry two protons across the membrane, as opposed to one, in the case of lactate or pyruvate. Therefore, succinate transport by MCTs is presumably more sensitive to ΔpH across the plasma membrane than for other MCT substrates. Succinate export from cells by MCTs is highly pH dependent, due to the requirement for intracellular succinate to exist as a monocarboxylate, and ΔpH driving proton release with succinate. However, in the case of succinate import into brown adipocytes, the intracellular concentration of succinate would be equivalent or higher than the extracellular concentration, which peaks in the hundreds of micromolar. How then is uptake and metabolism of micromolar concentrations of succinate achieved via MCTs, if uptake is operating against the succinate concentration gradient? One contributing factor is likely to be ΔpH across the plasma membrane. Since succinate uptake would bring two molar equivalents of protons, relative alkalinization of the cytosol could preferentially drive proton import using succinate as a co-transported species. This supposition is supported by the observation that interventions driving cytosolic alkalinization, i.e. adrenoreceptor agonism, promote succinate uptake into brown adipocytes. Indeed, the highly innervated nature of BAT and beige fat, coupled with the high capacity of these cells for succinate oxidation to support thermogenesis, may be a major reason why these cells avidly acquire extracellular succinate. The above parameters dictating succinate transport by MCTs also suggest that the metabolic state of BAT could be a major factor regulating succinate uptake in these cells. For example, it would be expected that intracellular acidification in brown adipocytes would promote succinate export, as opposed to import. Such local metabolic changes could also affect the capacity for BAT to handle systemic succinate.

Understanding the metabolic and environmental conditions that foster succinate uptake into BAT will offer us greater insight into how BAT regulates systemic health. Succinate transport into BAT promotes a range of physiological responses that are beneficial for systemic health. These include increased thermogenic activity, protection from metabolic disease, and decreased inflammation. Here we show both *in vivo* and *in vitro* that MCT proteins, particularly MCT1, have a capacity to sequester extracellular succinate. We demonstrate a pH dependency of succinate transport and show BAT preferentially transports protonated succinate. Our *in vivo* MCT1 KO data suggests a compensatory pathway for succinate uptake, presumably through MCT2 or another solute carrier protein. Future studies will need to be conducted to further elucidate the full repertoire transporters capable of transporting succinate into BAT.

## Author contributions

A.R and E.T.C. wrote the manuscript. S.W. conducted and designed all ^14^C_4_-succinate experiments. E.L.M, R.G, and S.W performed the *in vivo* tracing experiment following acute MCT inhibition using AZD3965. A.R. and N.T. conducted *in vivo* tracing experiment in MCT1 KO mice. H.X. performed proteomics to confirm MCT1 KO using BAT harvested from MCT1 KO mice. J.J. and R.G. conducted *in vitro* ^13^C_4_-succinate tracing experiment following treatment with AZD3965. A.R. performed imaging experiments to determine cytosolic pH. A.R. performed *in vitro* ^13^C_4_-succinate tracing experiment with NE. E.T.C, S.W., E.L.M, and A.R. oversaw the experiments and data analysis. All authors edited the manuscript.

## Acknowledgements

We thank Jeffrey Rothstein (Johns Hopkins) for providing the MCT1 flox mice. This work was supported by The Novo Nordisk Foundation Center for Basic Metabolic Research is an independent Research Center, based at the University of Copenhagen, and partially funded by an unconditional donation from the Novo Nordisk Foundation (Grant number NNF18CC0034900), a fellowship from the Novo Nordisk Foundation (NNF18OC0032380) (S.W), the Claudia Adams Barr Program (E.T.C), the Lavine Family Fund (E.T.C), the Pew Charitable Trust (E.T.C), NIH DK123095 (E.T.C), NIH AG071966 (E.T.C), The Smith Family Foundation (E.T.C), and the American Federation for Aging Research (E.T.C).

